# Optimizing light environment enables speed breeding in forage legumes: physiological limits and generation time reduction in *Medicago sativa* and *Medicago truncatula*

**DOI:** 10.64898/2026.01.20.700577

**Authors:** Andrés Berais-Rubio, Camila Couture, Melanie Rodríguez-Briosso, Santiago Signorelli

## Abstract

Climate change and increasing global demand for animal products are intensifying the need to accelerate genetic improvement of forage crops. Speed breeding (SB) has emerged as a powerful tool to shorten generation cycles; however, its application in perennial and autotetraploid forage legumes remains limited, particularly regarding reproductive performance and physiological constraints. Here, we optimized photoperiod, light intensity, and light quality to accelerate the life cycle of *Medicago sativa* (alfalfa) and its diploid relative *Medicago truncatula* under controlled conditions. We evaluated flowering, fruiting, seed harvest time, seed set, and germination across independent and combined SB treatments, and assessed photosynthetic performance to identify potential physiological trade-offs. Blue– red light supplementation, moderate-to-high irradiance (450 µmol m^-2^ s^-1^), and extended photoperiods significantly accelerated reproductive development in both species, although optimal combinations differed between the diploid and autotetraploid backgrounds. A combined SB regime (20/4 h photoperiod at 450 µmol m^-2^ s^-1^) reduced time to harvest by 17% in *M. sativa* and 28% in *M. truncatula*, while maintaining viable seed production. Chlorophyll fluorescence analysis revealed a higher photosynthetic plasticity in alfalfa compared with *M. truncatula*, indicating species-specific physiological limits to SB intensification. Our results establish practical SB conditions for alfalfa and an agronomically relevant *M. truncatula* genotype, providing an enabling platform to accelerate breeding cycles and trait evaluation in forage legumes.

## Introduction

Climate change and global population growth are threatening food security, making it urgent to enhance the productivity of agricultural systems, especially in fundamental crops for food production and sustainability (Samantara et al., 2022; Temesgen, 2022). The highly erratic climate conditions, rising global temperatures, long droughts, and irregular rainfall patterns are reducing yields and compromising forage and global food production (Putnam, 2021; Samac & Temple, 2021). Forage legumes are fundamental in sustainable agriculture because they enrich soils through biological nitrogen fixation while providing high nutritional value for livestock (Alexandratos & Bruinsma, 2012; Hickey et al., 2019; Hussain et al., 2018).

Alfalfa (*Medicago sativa*) is the most economically significant forage crop worldwide, and it is used for silage, hay, and pasture, which makes it essential for livestock feeding and soil health (Yi & Kole, 2021). However, genetic improvement in alfalfa has been slow due to its self-incompatibility and complex autotetraploid genome (Parajuli et al., 2021; Zheng et al., 2022). In contrast, its diploid relative, *Medicago truncatula* constitutes a genetic model system for research in legumes because of its fully sequenced genome and short life cycle (Lesins & Lesins, 1979; Suttie, 2012). Despite progress achieved using *M. truncatula*, translating these findings to alfalfa has been complicated, not only because of its autotetraploidy and self-incompatibility, but also because of its longer generation time, which slows down breeding cycles in the purchase of genetic improvements (Temesgen, 2022; Watson et al., 2018). Recent advances in genome editing technologies, such as CRISPR/Cas9, combined with the availability of chromosome-level genome assemblies for alfalfa, have opened new possibilities for efficient genetic manipulation and accelerated breeding in this crop (Zhao et al., 2024; Zheng et al., 2022). In this scenario, it is crucial to develop techniques that reduce alfalfa breeding time and improve its adaptability to environmental stresses (Potts et al., 2023).

One promising approach is Speed Breeding (SB), which optimizes environmental parameters such as photoperiod, temperature, humidity, and light quality to accelerate crop development and seed production (Ahmar et al., 2020; Bhatta et al., 2021; Bonea, 2022; Ghosh et al., 2018; H. Li et al., 2019; Samineni et al., 2020). The concept was popularized by Watson et al., (2018), who developed protocols enabling up to six generations per year in several crops, including *M. truncatula*, using a 22/2 h light/dark photoperiod and LED lighting enriched in blue, red, and far-red wavelengths at 360–380 µmol.m^-2^.s^-1^. SB has been successfully implemented in multiple crop species, including chickpea (*Cicer arietinum*), where up to seven generations per year were achieved through extended photoperiod and immature seed germination, and wheat (*Triticum aestivum*), where shuttle breeding systems combined with controlled environments have accelerated generation turnover in breeding programs (H. Li et al., 2019; Samineni et al., 2020). Recent studies have extended SB to warm-season legumes such as cowpea (*Vigna unguiculata*; Edet & Ishii, 2022) and temperate legumes such as faba and pea bean (*Vicia faba, Pisum sativum*; Cazzola et al., 2020; Mobini et al., 2020). In contrast, *M. sativa* has received limited attention. Only very recently, Han et al., (2025) reported combinations of light intensity and photoperiod that increased vegetative growth in *M. sativa*. However, this study did not evaluate key reproductive variables and thus does not align with the core objectives of SB. Consequently, the effects of light quality, intensity, and photoperiod on the reproductive performance of this autotetraploid species remain largely unexplored.

Physiologically, SB approaches must balance rapid development with optimal plant performance. Excessive light intensity or prolonged photoperiods can induce photoinhibition, oxidative stress, or imbalances in carbon partitioning, potentially reducing photosynthetic efficiency, biomass accumulation, and reproductive success (Bhatta et al., 2021). Chlorophyll fluorescence parameters are reliable indicators of photosynthetic performance and provide a convenient, non-invasive method to assess the functional state of the photosynthetic apparatus (Calzadilla et al., 2022). Moreover, changes in these parameters can indirectly reflect oxidative stress or photoinhibitory damage in response to excess light (Signorelli et al., 2013). Therefore, defining the upper physiological limits for SB parameters is essential, particularly in species differing in genome complexity and ploidy level, such as *M. truncatula* and *M. sativa*.

We hypothesize that by optimizing light intensity, photoperiod, and light quality, the generation time of *M. truncatula* and *M. sativa* can be shortened without negatively affecting seed set or germination. We further expect the optimal combination of parameters to differ between the diploid model and the autotetraploid crop species. The development of such SB protocols would facilitate the use of advanced breeding technologies (He & Li, 2020; Zhao et al., 2024), including gene editing and genomic selection, allowing breeders to generate and evaluate new lines more rapidly (Wolter et al., 2019; He & Li, 2020).

In this study, we evaluated the effect of light quality (Full-spectrum vs Blue-Red), light intensity (200, 450, and 650 µmol.m^-2^.s^-1^), and photoperiod (16/8, 20/4, and 22/2 light/dark hours) on the development of both *M. sativa* and *M. truncatula* under controlled conditions. We assessed flowering time, fruiting time, days to harvest, the number of seeds per fruit, and germination rate, and then tested a combined optimal condition integrating the best parameters for each species.

## Materials and methods

### Plant material and general growing conditions

Seeds of *M. sativa cv*. Chaná and *M. truncatula* CALIPH were superficially sterilized according to Irisarri et al. (2019). For each treatment, five pots were used, each containing five plants. Plants were irrigated alternately with sterile water and Fahraeus nutrient solution composed of 0.08 g CaCl_2 2_H_2_O, 0.06 g MgSO_4 7_H2O, 0.06 g NaCl, 0.2 g K_2_HPO_4_, 0.17 g K_2_H_2_PO_4_, 0.005 g ferric citrate, and 1 mL. L^-1^ of micronutrient solution (Vincent, 1970).

Control plants were grown at 23°C (day/night), under 16/8 h (light/dark) photoperiod, using Full-spectrum LED lamps (Nanolux, USA) providing a photon flux density (PFD) of 200 µmol.m^-2^.s^-1^ at bench height (280 µmol.m^-2^.s^-1^ at 50 cm above the pot). Light spectrum and intensities were verified using a USB2000+ spectroradiometer (Ocean Optics, Netherlands).

### Light quality assays

To evaluate the effect of light quality, plants were grown in a controlled room at 23°C and fluctuating relative humidity between 40 to 65%. Treatments included Full-spectrum light (control) and Blue-Red light (450nm + 660nm; Nanolux, USA), as is shown in Figure 1. Both treatments delivered a PFD of 200 µmol.m^-2^.s^-1^ at bench height (∼280 µmol.m^-2^.s^-1^ at 50 cm above the pot), under a 16/8 h photoperiod. Blue-Red bars were operated at 34% of the total power output.

**Figure 1.**
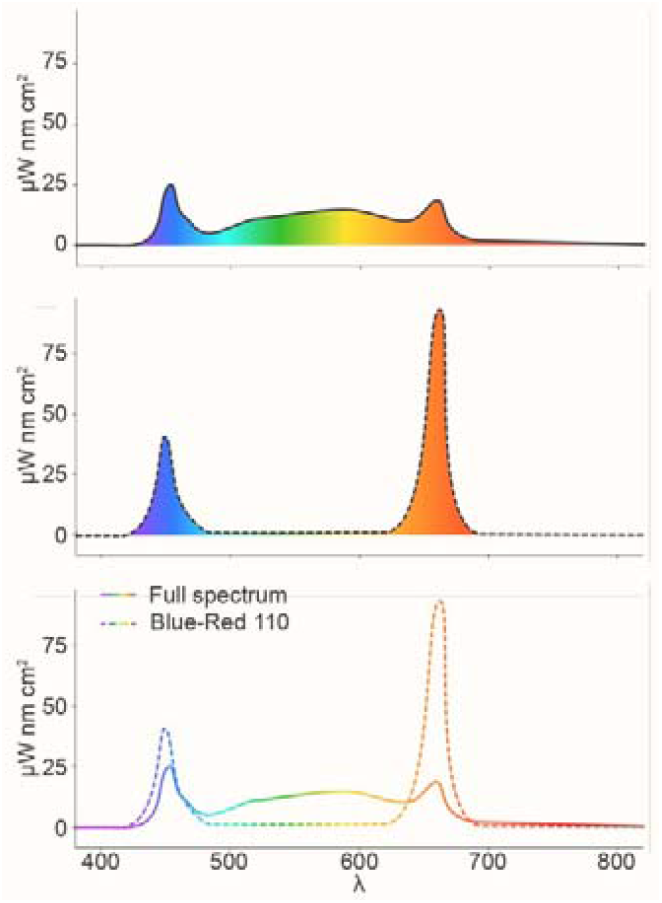
Comparison of light spectrum: Full-spectrum and Blue-Red. The solid line represents a Full-spectrum light, approximately 400-700 nm and beyond. The dashed line shows a Blue-Red spectrum, characterized by two distinct peaks corresponding to blue (∼450 nm) and red (∼660 nm) wavelengths. The y-axis shows spectral irradiance in microwatts per nanometer per square centimeter (μW.nm ^−1^ cm ^−2^), while the x-axis displays the wavelength (λ) in nanometers.

### Light intensity assays

To test the effect of light intensity, plants were grown in a Biora 800L RIC growth chamber (MineARC Systems, Australia) under Full-spectrum lighting (16/8 h photoperiod). PFD at bench height was adjusted to 200 µmol.m^-2^.s^-1^ (control), 450 µmol.m^-2^.s^-1^ (35% lamp power), and 650 µmol.m^-2^.s^-1^ (50% lamp power), corresponding to approximately 280, 550, and 750 µmol.m^-2^.s^-1^ at 50 cm above the pots. Temperature and relative humidity were maintained at 21 °C and 50%, respectively.

### Photoperiod conditions assays

Photoeriod effects were tested using the same growth chamber and Full-spectrum light (PFD = 200 µmol.m^-2^.s^-1^ at bench height). Photoperiods of 16/8 h (control), 20/4 h, and 22/2 h (light/dark) were evaluated under controlled conditions (21 °C, 50% relative humidity). Lamp power was set to 10% to maintain the target intensity.

### Combined optimal speed breeding conditions

Based on results from the individual trials, a combined “optimal” condition was tested for each species. Plants were grown at 23 °C and 40-65% relative humidity under a mix of two Full-spectrum LED bars and one blue–red bar (Nanolux, USA). For both *Medicagos*, the selected SB condition was 450 µmol.m^-2^.s^-1^ and a 20/4 h photoperiod.

### Evaluated development-related parameters

For each treatment, the following parameters were recorded: i) flowering time: days from sowing until the appearance of the first closed flower per pot; ii) fruiting time: days from sowing until the first visible fruit; iii) days to harvest: days from sowing until the first dry fruit was observed; iv) seeds per fruit: number of seeds from a pool of 20 mature fruits per treatment for *M. sativa*, and from quadruplicates of 5 pools of 5 mature fruits per treatment for *M. truncatula*; v) seed germination rate: percentage of seeds germinated under standard growth conditions, three replicates per treatment with ∼40 seeds per replicate.

### Photosynthetic performance under SB conditions

Photosynthetic performance was assessed by simultaneous measurements of chlorophyll a fluorescence and P700 absorbance using a DUAL-PAM-100 fluorometer (WALZ, Germany). These measurements allowed the estimation of photosystem II (PSII) and photosystem I (PSI) activity in plants grown under control and combined optimal speed breeding (SB) conditions.

The following parameters were recorded: maximum quantum yield of PSII (Fv/Fm) after dark adaptation; steady-state quantum yields of PSI and PSII (Y(I) and Y(II)); electron transport rates through PSI and PSII (ETRI and ETRII); non-photochemical quenching (NPQ); donor- and acceptor-side limitations of PSI (Y(ND) and Y(NA)); and photochemical quenching parameters of PSII (qP and qL).

For each treatment, five pots were used, each containing five plants. Measurements were performed on one fully expanded leaf per pot, with two technical measurements per plant. Leaves were dark-adapted for 20 min prior to measurements. Measuring light and saturating pulse intensities were set to 34 and 10,286 µmol.m^-2^.s^-1^, respectively for both species, while actinic light intensity was set to 894 and 722 µmol.m^-2^.s^-1^, for alfalfa and *M. truncatula* respectively.

### Statistical analysis

Statistics and plotting were performed using R 4.3.2 + ggplot2 (*R Core Team*, 2023; Wickham et al., 2023). ANOVA was performed in R software to identify statistical significance. A post hoc test (Tukey’s test) was used for pairwise comparison to determine the *p*-values.

### Use of AI-tools for manuscript preparation

Chat-GPT 5 and Elicit were employed to revise our initial draft and help us finding relevant literature. The AI-tools were selected considering they have shown to have at least 80% of accuracy in providing information related to our field of research (Fernandez Burda et al., 2025). All suggested information with its relevant citations were manually revised by the authors of this study.

## Results

To identify growth conditions that accelerate the life cycle of *M. sativa* and *M. truncatula*, we independently evaluated the effects of light quality, light intensity, and photoperiod under controlled-environment conditions. For each treatment, flowering time, fruiting time, and days to harvest were recorded, followed by the evaluation of seed set and germination rate.

### Effect of light quality on the life cycle of *M. sativa* and *M. truncatula*

In *M. sativa*, Blue–Red illumination significantly reduced flowering time (*p* < 0.001) and days to harvest (*p* < 0.05) compared with Full-spectrum light, while fruiting time remained unaffected (Figure 2). In *M. truncatula*, Blue-Red light accelerated all developmental stages: flowering (*p* < 0.01), fruit set (*p* < 0.05), and harvest (*p* < 0.05). Overall, Blue-Red lighting consistently promoted earlier reproductive transitions in both species, with stronger effects observed in *M. truncatula* (Figure 2).

**Figure 2.**
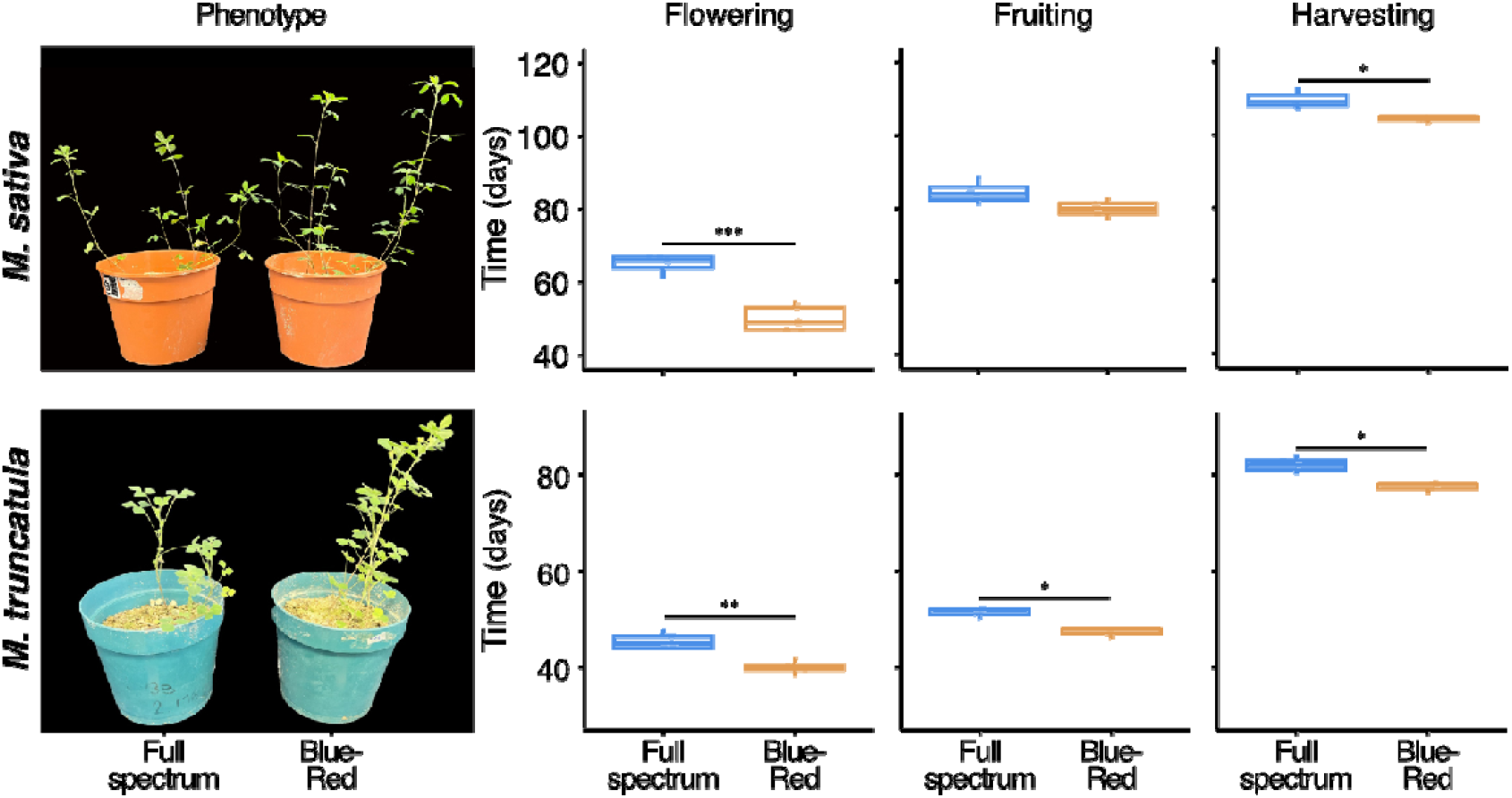
Effect of light quality on reproductive traits of *M. sativa* and *M. truncatula*. Plants of *M. sativa* cv. Chaná and *M. truncatula* CALIPH were grown under controlled environmental conditions to examine how different light spectra influence reproductive timing. Light quality treatments consisted of Full-spectrum LED lighting (control) and a blue–red LED spectrum (450 nm + 660 nm; Nanolux, USA). Both treatments delivered a photon flux density (PFD) of 200 μmol m□^2^ s□^1^ at bench height (∼280 μmol m□^2^ s□^1^ at 50 cm above pot level) under a 16/8 h light/dark photoperiod. Blue–red LED bars were operated at 34% of total power output, ensuring equivalent PFD between treatments. Growth room conditions were maintained at 23 °C with 40–65% relative humidity, and plants were irrigated with sterile water and Fahraeus nutrient solution on alternating days. For each treatment, five pots, each containing five plants, were monitored for key developmental transitions: (i) flowering time (days from sowing to appearance of the first closed flower), (ii) fruiting time (days to the first visible pod), and (iii) days to harvest (days until the first dry, harvestable pod). Plots display the distribution of pot-level measurements for each species, with grey points indicating individual observations. Statistical comparisons between light quality treatments were performed using ANOVA followed by Tukey’s post hoc test. Asterisks indicate significant differences (\**p* < 0.05, \*\**p <* 0.v01, \*\*\**p <* 0.001), while the absence of asterisks denotes non-significant contrasts. Error bars represent the standard error of the mean (SEM).

### Effect of light intensity on the life cycle of *M. sativa* and *M. truncatula*

Increasing light intensity markedly accelerated the reproductive cycle of *M. sativa*. Both 450 and 650 µmol.m^-2^.s^-1^ treatments significantly reduced flowering, fruiting, and harvest times compared with the control (Figure 3, *p* < 0.001 for all comparisons). Among these, 450 µmol.m^-2^.s^-1^ was the most effective, producing significantly earlier flowering and fruiting than 650 µmol.m^-2^.s^-1^ (Figure 3, *p* < 0.001 and *p* < 0.05, respectively).

**Figure 3.**
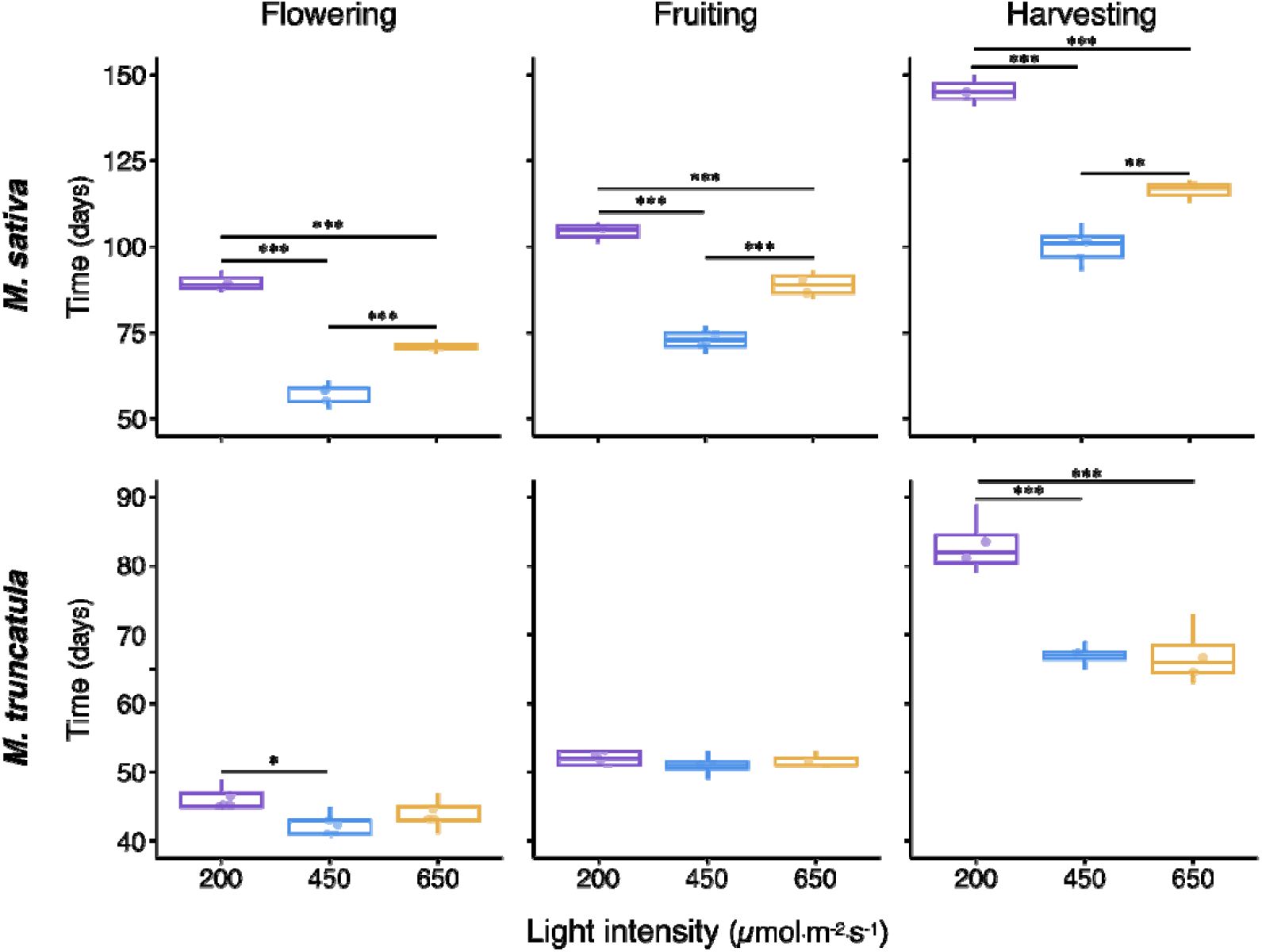
Effect of light intensity on reproductive traits of *M. sativa* and *M. truncatula*. Plants of *M. sativa* cv. Chaná and *M. truncatula* CALIPH were grown under controlled environmental conditions in a Biora 800L RIC growth chamber to evaluate the impact of different light intensities on key reproductive traits. Full-spectrum LED lighting (16/8 h light/dark photoperiod) was supplied at three photon flux densities (PFD): 200 μmol m□^2^ s□^1^ (control), 450 μmol m□^2^ s□^1^ (35% lamp power), and 650 μmol m□^2^ s□^1^ (50% lamp power), corresponding to approximately 280, 550, and 750 μmol m□^2^ s□^1^ at 50 cm above the pots, respectively. Temperature and relative humidity were maintained at 21 °C and 50%, and plants were irrigated alternately with sterile water and Fahraeus nutrient solution. For each intensity treatment, five pots per species, each containing five plants, were monitored throughout their life cycle. The reproductive parameters quantified were: (i) flowering time (days from sowing to appearance of the first closed flower); (ii) fruiting time (days to first visible pod); and (iii) days to harvest (days until the first dry, harvest-ready pod was observed). Boxes represent the distribution of measurements per treatment, and grey points indicate individual pot values. Statistical differences among light intensities were determined using ANOVA followed by Tukey’s post hoc test. Asterisks denote significant pairwise differences between treatments (\**p* < 0.05, \*\**p* < 0.01, *** *p* < 0.001), while absence of asterisks indicates non-significant contrasts. Error bars indicate the standard error of the mean (SEM).

In *M. truncatula*, the 450 µmol.m^-2^.s^-1^ treatment also accelerated flowering relative to the control (Figure 3, *p* < 0.05). However, no significant differences were observed between the two higher intensities (450 and 650 µmol.m^-2^.s^-1^). Days to harvest were reduced under both SB intensities (Figure 3, *p* < 0.001), indicating that moderate-to-high irradiance promotes faster development without major differences between these levels.

### Effect of photoperiod on reproductive transitions of *M. sativa* and *M. truncatula*

The photoperiod length significantly influenced the timing of reproductive events. In *M. sativa*, plants grown under a 20/4 h light/dark regime flowered significantly earlier than those under 16/8 h or 22/2 h conditions (Figure 4, *p* < 0.01). Fruiting time was not affected, but harvest occurred earlier under 20/4 h (Figure 4, *p* < 0.01).

**Figure 4.**
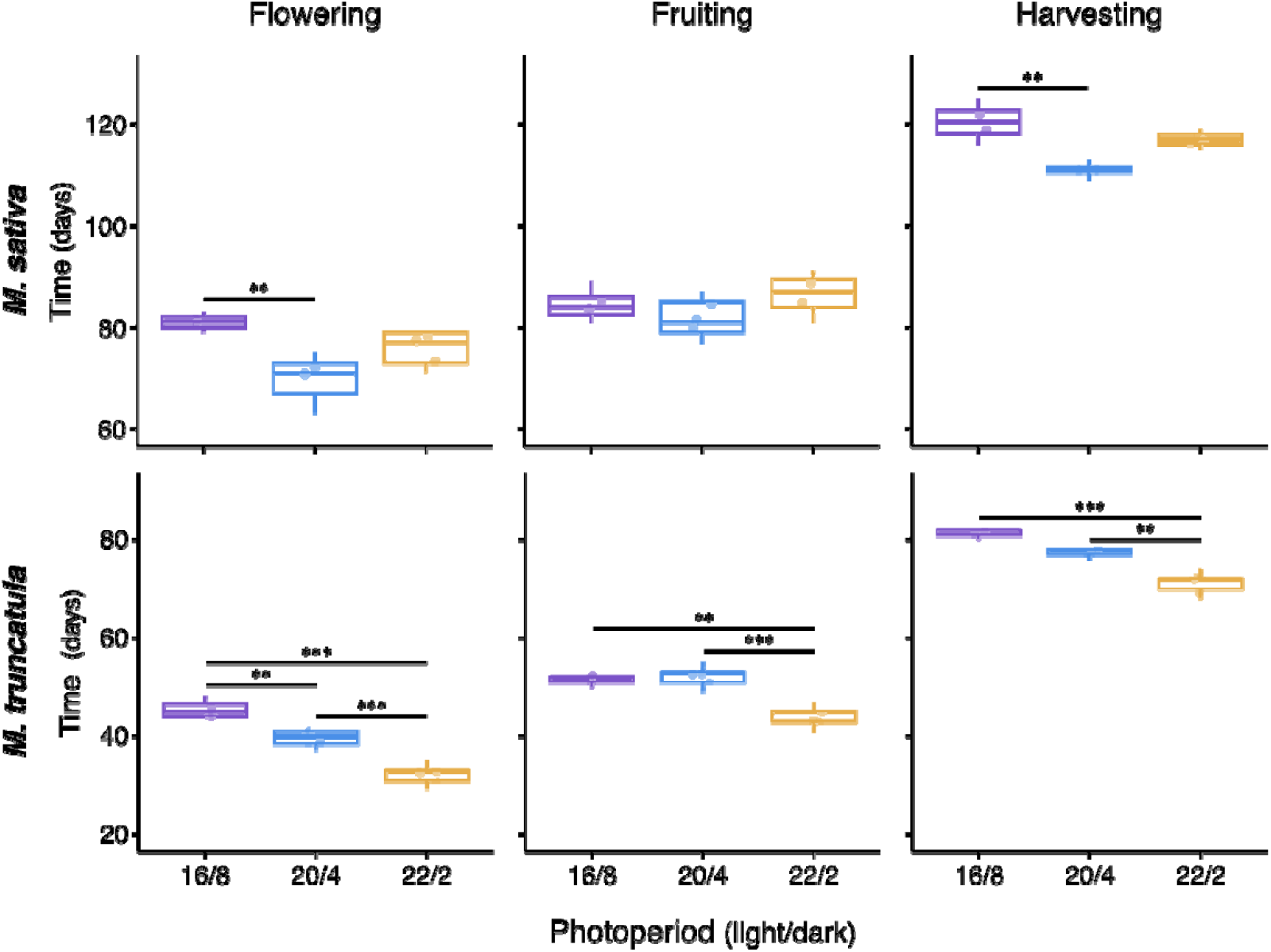
Effect of photoperiod on reproductive traits of *M. sativa* and *M. truncatula*. To assess the influence of day length on reproductive timing, *M. sativa* cv. Chaná and *M. truncatula* CALIPH were grown under controlled environmental conditions in a Biora 800L RIC growth chamber. Plants were exposed to three photoperiod regimes: 16/8 h, 20/4 h, and 22/2 h (light/dark). All treatments were performed under Full-spectrum LED lighting delivering a photon flux density of 200 μmol m□^2^ s□^1^ at bench height, corresponding to approximately 280 m□^2^ s□^1^ at 50 cm above the pots, maintained by operating the lamps at 10% power across all photoperiods. Growth conditions were kept stable at 21 °C and 50% relative humidity, and irrigation alternated between sterile water and Fahraeus nutrient solution. For each photoperiod treatment, five pots per species, each containing five plants, were monitored for key developmental stages: (i) flowering time (days from sowing to the appearance of the first closed flower), (ii) fruiting time (days until the first visible pod), and (iii) days to harvest (days until the first dry, mature pod was observed). Boxplots summarize pot-level data, with grey circles indicating individual observations. Statistical analyses were conducted using ANOVA followed by Tukey’s post hoc test to assess differences among photoperiod treatments. Asterisks denote significant pairwise differences (\**p* < 0.05, \*\**p* < 0.01, *** *p <* 0.001), while the absence of asterisks indicates non-significant contrasts. Error bars represent the standard error of the mean (SEM).

In contrast, *M. truncatula* responded more strongly to very long photoperiods. The 22/2 h and 20/4 h treatment conditions significantly accelerated flowering, while fruiting and harvesting were achieved earlier in 22/2 h treatment compared with 16/8 h and 20/4 h conditions (Figure 4).

### Seeds set and germination under each speed breeding condition

In *M. sativa*, the number of seeds per fruit did not present significant differences between treatments, except for the 22/2 treatment, which produced significantly more seeds than the Blue-Red treatment (Figure 5, *p* < 0.05). For the germination rate, the control treatment exhibited a significantly higher percentage of germinated seeds than the 20/4 h treatment (*p* < 0.05), 22/2 h treatment (*p* < 0.05), and 450 µmol.m^-2^.s^-1^ (*p* < 0.01) treatments (Figure 5). The 450 µmol.m^-2^.s^-1^ treatment also h ad a significantly lower germination rate (*p* < 0.05) than the 650 µmol.m^-2^.s^-1^ treatment.

**Figure 5.**
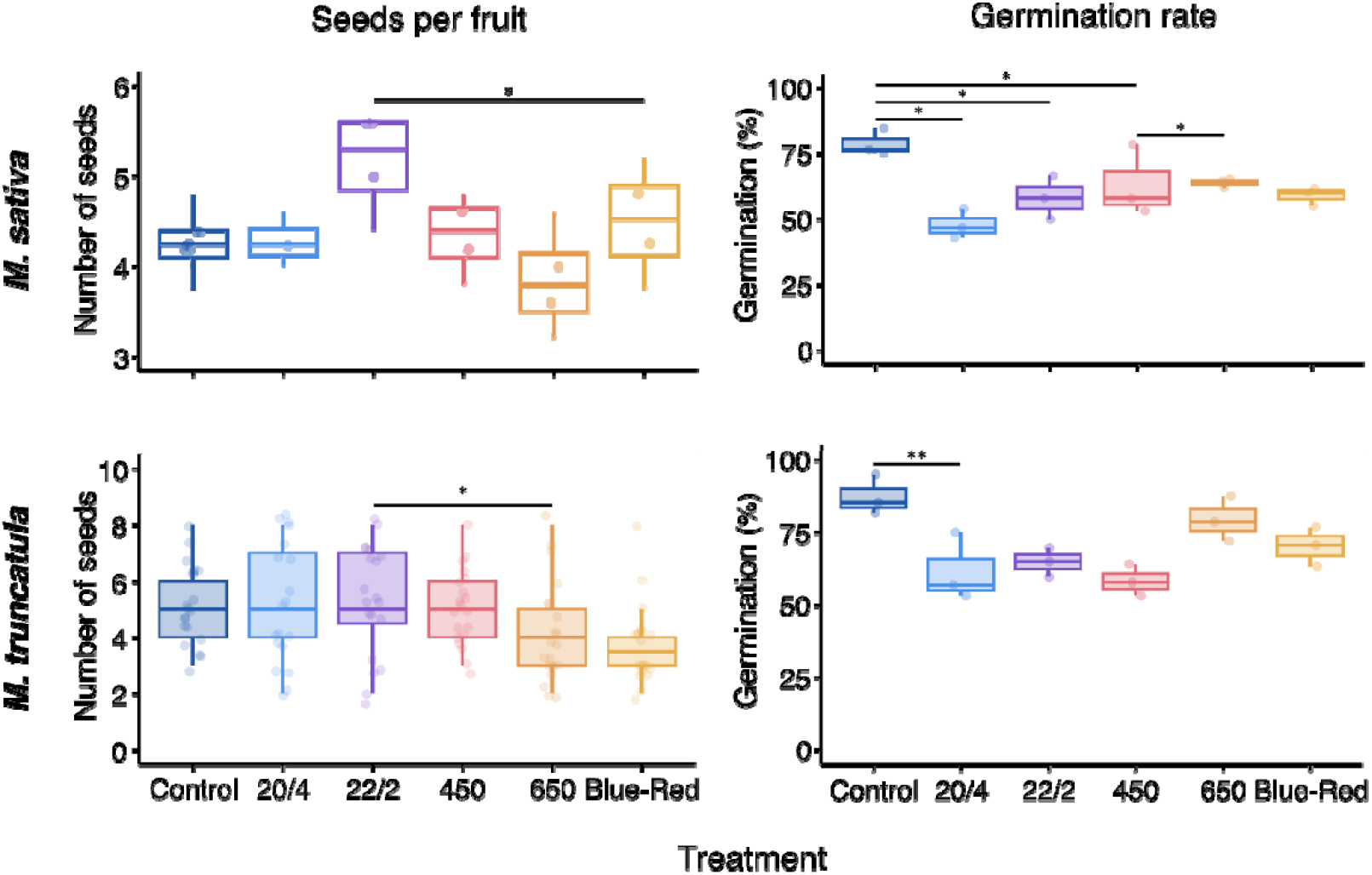
Effect of speed-breeding light treatments on seed set and seed germination in *M. sativa* cv. Chaná and *M. truncatula* CALIPH. A. Seeds per fruit measured for each light treatment and control. Treatments (x-axis) are: Control = Full-spectrum LED, 16/8 h (light/dark), PFD 200 µmol·m□^2^·s□^1^ at bench height; 20/4 and 22/2 = extended photoperiods (20 h light/4 h dark and 22 h light/2 h dark, respectively) under Full-spectrum LED; 450 and 650 = increased light intensities under Full-spectrum LED (PFD 450 and 650 µmol·m□^2^·s□^1^ at bench height, corresponding to 550 and 750 µmol·m□^2^·s□^1^ at 50 cm above the pot); Blue–Red = combined blue (450 nm) + red (660 nm) LED bars (operated at ∼34% total power) providing a PFD ≈ 200 µmol·m□^2^·s□^1^ at bench height. For each treatment, five pots were grown, each containing five plants. Seeds per fruit were scored from mature fruits as follows: *M. sativa*, a pooled sample of 20 mature fruits per treatment; *M. truncatula*, quadruplicate measurements consisting of 5 pools of 5 mature fruits per treatment (see Methods for full sampling details). Error bars indicate the standard error of the mean. B. Seed germination percentage for seeds collected under each light treatment and germinated under standard growth conditions. Germination percentage was calculated as the proportion of seeds showing radicle emergence during the germination assay. For both plots, an analysis of variance (ANOVA) was performed on pooled biological replicates, followed by Tukey’s honestly significant difference (HSD) post-hoc test for pairwise comparisons. Asterisks denote significant pairwise differences (\**p* < 0.05, \*\**p* < 0.01), while the absence of asterisks indicates non-significant contrasts. Error bars represent the standard error of the mean (SEM).

For *M. truncatula*, the number of seeds per fruit did not differ significantly between control and SB treatments, except under the 22/2 h photoperiod and 650 µmol.m^-2^.s^-1^ intensity, where fruit set was slightly reduced (Figure 5A), possibly due to stress at high irradiance. Germination percentage was similar across treatments, except for the 20/4 h photoperiod, in which seeds from control plants showed a slightly higher germination rate (Figure 5B, *p* < 0.05).

### Combined optimal speed breeding conditions

After defining the most effective parameters for each variable, we tested a combined condition for both species under a mix of two Full-spectrum LED bars and one Blue-Red bar (Nanolux, USA).: 20/4 h photoperiod at 450 µmol.m^-2^.s^-1^ (Figure 6). Firstly, we evaluated the combined condition of 22/2 h at 650 µmol.m^-2^.s^-1^ for *M. truncatula*, but the stacking of these demanding photoperiod and light intensity was deleterious for the plants (Figure 8). They could not develop properly, and therefore we explored the same conditions for both *Medicago* which implicated a reduced light intensity and a reduction of two hours in terms of photoperiod.

**Figure 6.**
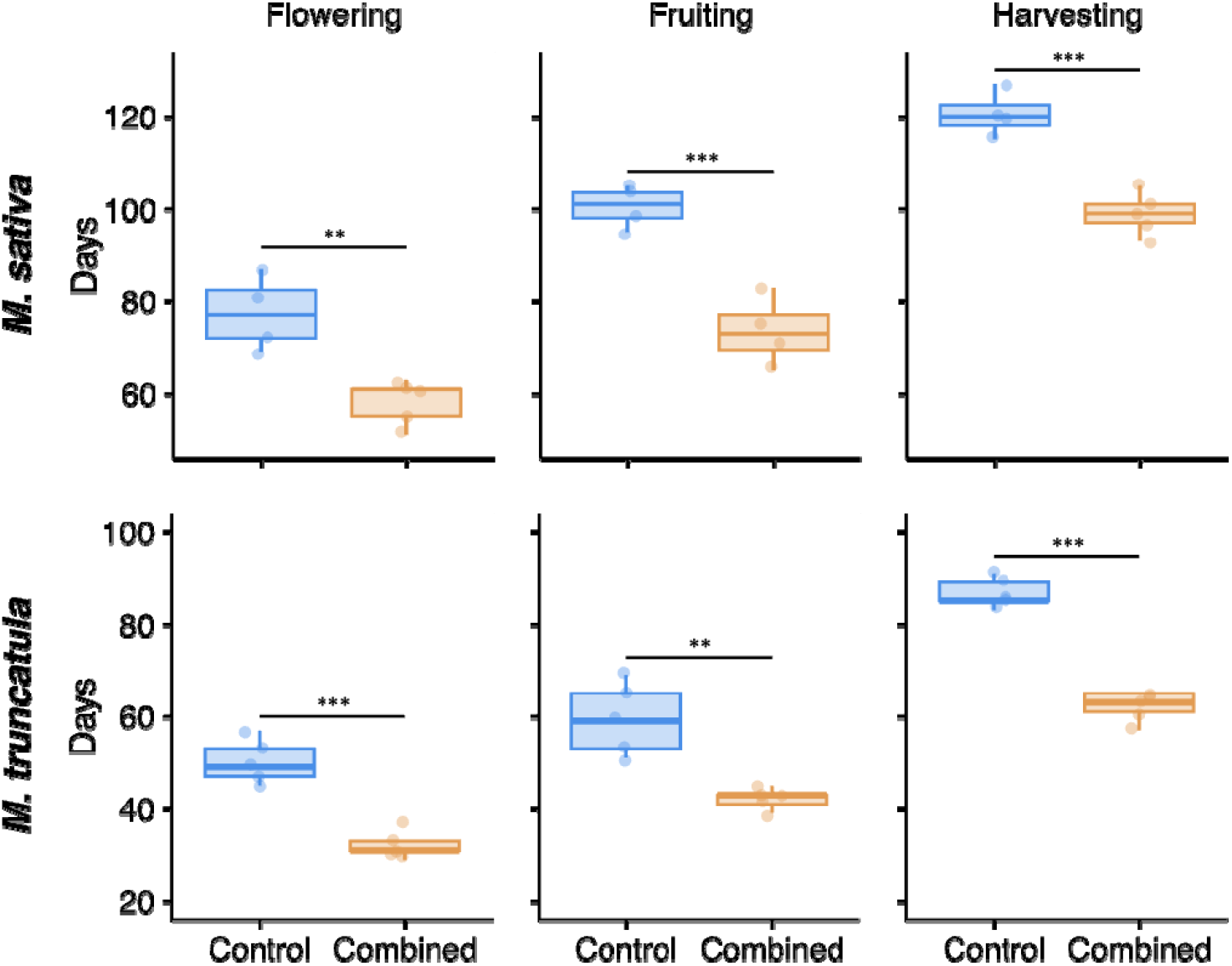
Effect of combined speed breeding conditions on reproductive traits of *M. sativa* and *M. truncatula*. To assess the effect of the combined conditions of light quality, intensity, and photoperiod, *M. sativa* cv. Chaná and *M. truncatula* CALIPH were grown under controlled environmental conditions at 23 ºC. In combined conditions, plants were exposed to a photoperiod of 20/4 h, under a mix of two Full-spectrum LED bars and one blue–red bar (Nanolux, USA), and light was supplied at the photon flux densities (PFD) of 450 μmol m□^2^ s□^1^ (35% lamp power), corresponding to approximately 550 μmol m□^2^ s□^1^ at 50 cm above the pots, respectively. Control conditions were performed under Full-spectrum LED lighting delivering a photon flux density of 200 μmol m□^2^ s□^1^ at bench height, corresponding to approximately 280 m□^2^ s□^1^ at 50 cm above the pots, maintained by operating the lamps at 10% power across all photoperiods, and a photoperiod of 16/8 h. Irrigation alternated between sterile water and Fahraeus nutrient solution. For each treatment, five pots per species, each containing five plants, were monitored for key developmental stages: (i) flowering time (days from sowing to the appearance of the first closed flower), (ii) fruiting time (days until the first visible pod), and (iii) days to harvest (days until the first dry, mature pod was observed). Boxplots summarize pot-level data, with grey circles indicating individual observations. Statistical analyses were conducted using ANOVA followed by Tukey’s post hoc test to assess differences among photoperiod treatments. Asterisks denote significant pairwise differences (*p* < 0.01, *** *p <* 0.001), while the absence of asterisks indicates non-significant contrasts. Error bars represent the standard error of the mean (SEM).

Preliminary observations indicate that these combined SB conditions strongly reduced vegetative growth, resulting in compact plants that flowered earlier but with lower vigor.

With respect to the quantitative results on flowering, fruiting, and harvesting time under this combined treatment, both *Medicago* species achieved all these parameters significantly earlier than the control conditions (Figure 6), achieving a significant acceleration of harvesting days of 17 and 28 % for *M. sativa* and *M. truncatula*, respectively.

Regarding the number of seeds per fruit and germination rate in combined conditions, for *M. sativa* there were no statistical differences in seed set, but the control condition had a significantly higher germination rate than the combined condition (Figure 7, *p* < 0.05). For *M. truncatula*, no statistical differences were observed for seeds per fruit and germination rate (Figure 7).

**Figure 7.**
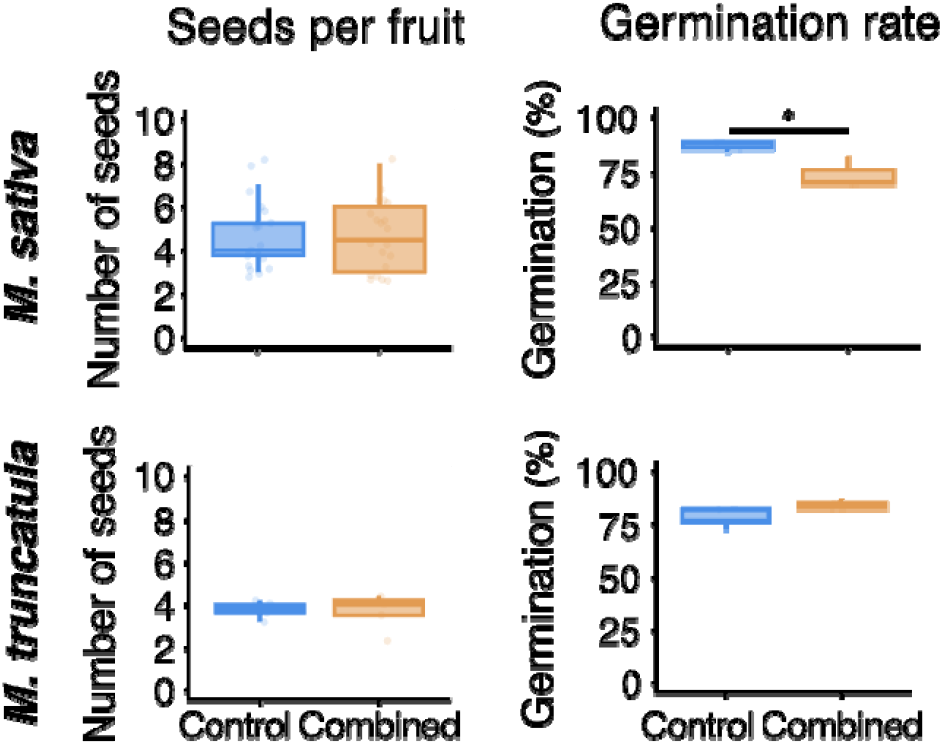
Effect of combined optimal conditions on seed set and seed germination in *M. sativa* cv. Chaná and *M. truncatula* CALIPH. A. Seeds per fruit measured for combined conditions treatment and control. In combined conditions, plants were exposed to a photoperiod of 20/4 h, under a mix of two Full-spectrum LED bars and one blue–red bar (Nanolux, USA), and light was supplied at the photon flux densities (PFD) of 450 μmol m□^2^ s□^1^ (35% lamp power), corresponding to approximately 550 μmol m□^2^ s□^1^ at 50 cm above the pots, respectively. Control conditions were performed under Full-spectrum LED lighting delivering a photon flux density of 200 μmol m□^2^ s□^1^ at bench height, corresponding to approximately 280 m□^2^ s□^1^ at 50 cm above the pots, maintained by operating the lamps at 10% power across all photoperiods, and a photoperiod of 16/8 h. Irrigation alternated between sterile water and Fahraeus nutrient solution. For each treatment, five pots were grown, each containing five plants. Seeds per fruit were scored from mature fruits as follows: *M. sativa*, a pooled sample of 20 mature fruits per treatment; *M. truncatula*, quadruplicate measurements consisting of 5 pools of 5 mature fruits per treatment (see Methods for full sampling details). Error bars indicate the standard error of the mean. B. Seed germination percentage for seeds collected under each treatment and germinated under standard growth conditions. Germination percentage was calculated as the proportion of seeds showing radicle emergence during the germination assay. For both plots, an analysis of variance (ANOVA) was performed on pooled biological replicates, followed by Tukey’s honestly significant difference (HSD) post-hoc test for pairwise comparisons. Asterisks denote significant pairwise differences (\**p* < 0.05), while the absence of asterisks indicates non-significant contrasts. Error bars represent the standard error of the mean (SEM).

### Photosynthetic performance under speed breeding conditions

To determine whether accelerated development under SB conditions was associated with changers in photosynthetic performance, chlorophyll fluorescence parameters related to PSII activity (Fv/Fm, ΦPSII [Y(II)], NPQ, qP) and P700 absorption-derived parameters describing PSI function (ΦPSI [Y(I)], Y(ND), Y(NA)) were evaluated in plants grown under control and combined SB conditions.

In *M. sativa*, none of the analysed parameters differed significantly between control and SB treatments (Figure 8). Maximum quantum yield of PSII (Fv/Fm), effective quantum yield of PSII (Y(II)), non-photochemical quenching (NPQ), photochemical quenching (qP), and PSI-related parameters (Y(I), Y(ND), and Y(NA)) remained unchanged, indicating the absence of photodamage and suggesting a high photosynthetic plasticity of alfalfa under intensified light regimes.

**Figure 8.**
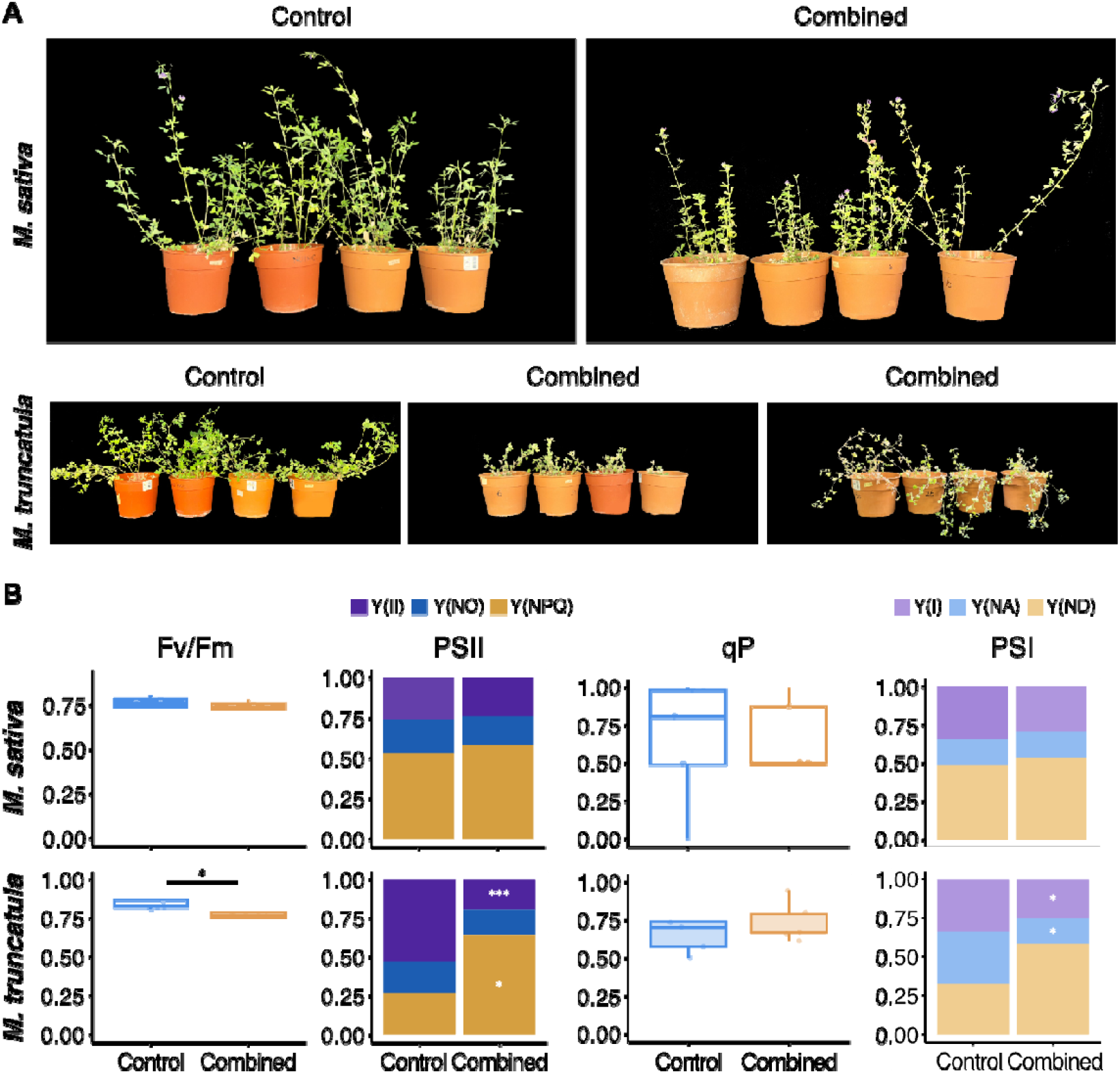
Photosynthetic performance under control and combined optimal conditions *M. sativa* cv. Chaná and *M. truncatula* CALIPH. Plants were grown under control conditions (23°C, under Full-spectrum LED lighting, 16/8 h light/dark photoperiod, 200 µmol·m ^2^·s ^1^) and combined optimal conditions (23°C, under a mix of two Full-spectrum LED bars and one blue–red bar, 20/4 h light/dark photoperiod, 450 µmol·m ^2^·s ^1^). Photosystem II (PSII) and photosystem I (PSI) parameters were assessed using a DUAL-PAM fluorometer. The displayed parameters include the maximum quantum yield of PSII (Fv/Fm), the effective quantum yield of PSII Y(II), the photochemical quenching coefficient of PSII (qP), and the effective quantum yield of PSI Y(I). Statistical analysis was performed using ANOVA followed by Tukey’s post hoc test for pairwise comparisons, with data pooled from three independent assays (each n=5), resulting in 12 total replicates per treatment. Bars represent means ± SE, and different letters indicate statistically significant differences (*p* < 0.05, *p* < 0.001) between treatments, and no asterisks indicate non-significant differences.

In contrast, *M. truncatula*, exhibited significant alterations in photosynthetic performance under SB conditions. At the level of PSII, plants showed reduced maximum quantum yield of PSII (Fv/Fm) and effective quantum yield of PSII Y(II), together with a marked increase in non-photochemical quenching (NPQ) (Figure 8). These changes indicate a reduced photochemical efficiency of PSII accompanied by enhanced energy dissipation mechanisms. Consistently, PSI activity was also affected, as evidenced by a reduction in the quantum yield of PSI (Y(I)) and a lower acceptor-side limitation (Y(NA)) in SB-grown plants (Figure 8), suggesting that decreased PSII activity constrains electron flow to PSI under these conditions.

## Discussion

In the context of achieving a sustainable agriculture, forage legumes are very important crops due to their capacity to substitute a significant portion of chemical nitrogen fertilizers while maintaining crop yields (Wei et al., 2024). However, legume productivity faces major challenges in acidic soils, where phosphorus deficiency and aluminum toxicity severely affect crop growth and yield, while drought stress remains a critical environmental factor limiting forage legume production worldwide (X. Li et al., 2023; Luo et al., 2023; Wei et al., 2024). Thus, breeding programs are still needed to develop new cultivars with higher resilience to these adversities.

SB has emerged as a powerful strategy to accelerate crop improvement by shortening generation cycles while maintaining reproductive competence and plant performance. Although SB protocols have been successfully implemented in several annual crops and model species, their application to forage legumes, particularly perennial and autotetraploid crops such as alfalfa, remains limited. In this study, we demonstrate that the generation time of *M. sativa* can be significantly reduced through optimized manipulation of the light environment without compromising seed viability, and that optimal SB configurations differ markedly from those required for its diploid relative *M. truncatula*. These results highlight the need for species- and genotype-specific SB strategies in forage crop improvement.

By independently dissecting the effects of photoperiod, light intensity, and light quality, and subsequently evaluating a combined SB regime, our results show that SB efficiency is not solely determined by maximal acceleration of developmental transitions, but is constrained by species-specific physiological limits. This distinction is particularly relevant for breeding programs, where excessive intensification of SB conditions may reduce plant vigor or seed quality, ultimately offsetting the benefits of shorter breeding cycles. Our findings therefore, support the concept that effective SB protocols must balance developmental acceleration with physiological stability, especially in crops with complex genomes such as autotetraploid alfalfa.

Consistent with previous SB studies, flowering was the developmental stage most responsive to SB conditions in both species (Watson et al., 2018). However, acceleration of flowering did not translate proportionally into shorter times to harvest, indicating that later reproductive stages impose additional constraints on overall generation time. By explicitly quantifying fruiting time, an aspect not evaluated in earlier SB studies on *Medicago*, we provide new insight into how early acceleration of anthesis may partially decelerate during pod development. This decoupling between flowering and harvest time underscores the importance of evaluating complete reproductive cycles when designing SB protocols for breeding purposes.

The optimal SB parameters differed between *M. sativa* and *M. truncatula*, reinforcing the notion that SB responses are strongly influenced by species biology and genome organization. In alfalfa, a 20/4 h photoperiod combined with moderate irradiance (450 µmol.m^-2^.s^-1^) resulted in the most consistent acceleration of harvesting time, whereas *M. truncatula* responded more strongly to extended photoperiods and higher light intensities, as previously reported for the Jemalong A17 reference line (Watson et al., 2018). Importantly, our study used the cultivar CALIPH, an agronomically relevant *M. truncatula* genotype developed for pasture systems, which contrasts with the laboratory-focused reference line A17. Differences in SB responses between these genotypes highlight the importance of evaluating SB protocols using material representative of breeding programs rather than exclusively model accessions. Given that several *M. truncatula* cultivars (e.g. CALIPH, Paraggio, Mogul) are used as annual forage legumes in Mediterranean-type environments, particularly in Australia and North Africa, understanding genotype-specific SB responses has direct relevance for forage improvement.

Across SB treatments, both species produced viable seeds, although seed number per fruit and germination rates varied among conditions. In alfalfa, seed production and germination remained within agronomically acceptable ranges across all SB regimes, indicating that accelerated cycles can be achieved without major penalties on reproductive output. In *M. truncatula*, seed viability was similarly maintained, although extreme SB conditions tended to reduce reproductive performance. These results reinforce the need to avoid overly aggressive SB regimes when the goal is sustained multi-generation advancement rather than rapid phenotyping alone.

Physiological assessments further revealed contrasting responses between the two species. Alfalfa plants grown under combined SB conditions showed no evidence of photodamage, maintaining stable PSII and PSI efficiencies relative to control plants. This indicates a high degree of photosynthetic plasticity and suggests that alfalfa can tolerate intensified light environments without compromising photosynthetic function. Conversely, *M. truncatula* displayed some level of photodamage under the combined optimal condition, evidenced by reduced Fv/Fm levels, a dark-adapted parameter. A direct comparison between the steady state parameters cannot be taken as the intensity of actinic light used were not the same. Our results, indicated *M. truncatula* exhibited reduced effective PSII efficiency under SB conditions, accompanied by increased non-photochemical quenching. These responses are indicative of photoinhibitory stress and enhanced energy dissipation, consistent with reports of light stress responses in other legumes such as pea, common bean, and soybean (Harbinson et al., 1989, Narina et al., 2014; Sánchez-Reinoso et al., 2019). Together, these results suggest that physiological limits to SB intensification are reached earlier in *M. truncatula* than in alfalfa, emphasizing the importance of tailoring SB regimes to species-specific tolerance thresholds.

Previous SB studies in alfalfa have primarily focused on vegetative growth or flowering responses under modified photoperiods, nutrient regimes, or light spectra, without addressing full reproductive cycles or physiological constraints (Chen et al., 2023, Han et al., 2025,Sysoeva et al., 2010). Our study extends this work by defining, for the first time, an integrated SB framework for alfalfa that combines optimized light quality, intensity, and photoperiod with evaluation of fruiting, seed viability, and photosynthetic performance. Similarly, for *M. truncatula*, we establish SB conditions for an agronomically relevant cultivar, moving beyond reference genotypes commonly used in controlled-environment research.

In practical terms, a 20/4 h photoperiod at 450 µmol.m^-2^.s^-1^ using a combination of full-spectrum and blue–red light represents an effective compromise for accelerating generation time in both *M. sativa* and *M. truncatula*, enabling seed harvest within approximately 100 and 62 days, respectively. Although greater acceleration has been reported in some annual crops such as pepper and pea (Liu et al., 2022; S. H. Mobini & Warkentin, 2016), the levels achieved here are substantial given the perennial nature and genome complexity of alfalfa. Collectively, our results provide a robust SB platform for forage legumes that can facilitate faster breeding cycles, trait evaluation, and integration with modern crop improvement tools such as genomic selection and gene editing.

## Conclusions

In this study, we successfully optimized speed breeding protocols for both *M. sativa* and *M. truncatula* under controlled environmental conditions. A combined speed breeding regime of 20/4 h photoperiod at µmol.m^-2^.s^-1^ with blue-red light supplementation achieved 17% and 28% reduction in time to harvest for alfalfa and *M. truncatula*, respectively, while maintaining viable seed production. Notably, alfalfa exhibited greater photosynthetic plasticity and tolerance to intensified light regimes compared to *M. truncatula*, which showed physiological stress under the same conditions, indicating species-specific physiological limits to speed breeding intensification. These results establish practical speed breeding frameworks for forage legumes that balance developmental acceleration with physiological stability, providing an enabling platform to accelerate breeding cycles and facilitate the integration of advanced breeding technologies in sustainable agriculture.

## Author contributions

Conceptualization: SS; Formal analysis: AB, CC, and SS; Funding acquisition: SS; Investigation: AB, CC, and MR; Methodology: AB and SS; Project administration: AB and SS; Resources: SS; Supervision: SS; Visualization: all authors; Writing – original draft: AB; Writing – review and editing: AB and SS.

## Acknowledgements

We thank Federico Condon providing the *M. truncatula* seeds used in this research, Dr. Omar Borsani for giving advise on how to cross *Medicago* plants, Dr. Pablo Calzadilla for contributing to setting up protocols for DUAL PAM analysis. SS is a thankful active member of the National System of Researchers (SNI, *Sistema Nacional de Investigadores*) and PEDECIBA from Uruguay.

## References

Ahmar, S., Gill, R. A., Jung, K.-H., Faheem, A., Qasim, M. U., Mubeen, M., & Zhou, W. (2020). Conventional and Molecular Techniques from Simple Breeding to Speed Breeding in Crop Plants: Recent Advances and Future Outlook. International Journal of Molecular Sciences, 21(7), 2590. 10.3390/ijms21072590

Alexandratos, N., & Bruinsma, J. (2012). World Agriculture towards 2030/2050: the 2012 revision. https://www.fao.org/economic/esa

Bhatta, M., Sandro, P., Smith, M. R., Delaney, O., Voss-Fels, K. P., Gutierrez, L., & Hickey, L. T. (2021). Need for speed: manipulating plant growth to accelerate breeding cycles. Current Opinion in Plant Biology, 60, 101986. 10.1016/j.pbi.2020.101986

Bonea, D. (2022). SPEED BREEDING AND ITS IMPORTANCE FOR THE IMPROVEMENT OF AGRICULTURAL CROPS. “Annals of the University of Craiova - Agriculture Montanology Cadastre Series “, 52(1), 59–66. 10.52846/aamc.v52i1.1314

Calzadilla, P. I., Carvalho, F. E. L., Gomez, R., Lima Neto, M. C., & Signorelli, S. (2022). Assessing photosynthesis in plant systems: A cornerstone to aid in the selection of resistant and productive crops. Environmental and Experimental Botany, 201, 104950. 10.1016/j.envexpbot.2022.104950

Cazzola, F., Bermejo, C. J., Guindon, M. F., & Cointry, E. (2020). Speed breeding in pea (Pisum sativum L.), an efficient and simple system to accelerate breeding programs. Euphytica, 216(11), 178. 10.1007/s10681-020-02715-6

Chen, Y., Liu, J., & Liu, W. (2023). Enhancing growth, quality, and metabolism of nitrogen of alfalfa (Medicago sativa L.) by high red–blue light intensity. Journal of Plant Nutrition and Soil Science, 186(6), 661–672. 10.1002/jpln.202300216

Edet, O. U., & Ishii, T. (2022). Cowpea speed breeding using regulated growth chamber conditions and seeds of oven-dried immature pods potentially accommodates eight generations per year. Plant Methods, 18(1), 106. 10.1186/s13007-022-00938-3

Fernandez Burda, M., Ferrero, L., Gaggion, N., Fonouni-Farde, C., Crespi, M., Ariel, F., & Ferrante, E. (2025). What Large Language Models Know About Plant Molecular Biology. 10.1101/2025.08.31.672925

Ghosh, S., Watson, A., Gonzalez-Navarro, O. E., Ramirez-Gonzalez, R. H., Yanes, L., Mendoza-Suárez, M., Simmonds, J., Wells, R., Rayner, T., Green, P., Hafeez, A., Hayta, S., Melton, R. E., Steed, A., Sarkar, A., Carter, J., Perkins, L., Lord, J., Tester, M., … Hickey, L. T. (2018). Speed breeding in growth chambers and glasshouses for crop breeding and model plant research. Nature Protocols, 13(12), 2944–2963. 10.1038/s41596-018-0072-z

Han, L., Lv, Y., Zhang, Y., Zhao, X., Gao, P., Liang, Y., & Li, B. (2025). Optimizing the Light Intensity, Nutrient Solution, and Photoperiod for Speed Breeding of Alfalfa (Medicago sativa L.) Under Full-Spectrum LED Light. Agronomy, 15(9), 2067. 10.3390/agronomy15092067

Harbinson, J., Genty, B., & Baker, N. R. (1989). Relationship between the Quantum Efficiencies of Photosystems I and II in Pea Leaves. Plant Physiology, 90(3), 1029–1034. 10.1104/pp.90.3.1029

He, T., & Li, C. (2020). Harness the power of genomic selection and the potential of germplasm in crop breeding for global food security in the era with rapid climate change. The Crop Journal, 8(5), 688–700. 10.1016/j.cj.2020.04.005

Hickey, L. T., N. Hafeez, A., Robinson, H., Jackson, S. A., Leal-Bertioli, S. C. M., Tester, M., Gao, C., Godwin, I. D., Hayes, B. J., & Wulff, B. B. H. (2019). Breeding crops to feed 10 billion. Nature Biotechnology, 37(7), 744–754. 10.1038/s41587-019-0152-9

Hussain, H. A., Hussain, S., Khaliq, A., Ashraf, U., Anjum, S. A., Men, S., & Wang, L. (2018). Chilling and Drought Stresses in Crop Plants: Implications, Cross Talk, and Potential Management Opportunities. Frontiers in Plant Science, 9. 10.3389/fpls.2018.00393

Irisarri, P., Cardozo, G., Tartaglia, C., Reyno, R., Gutiérrez, P., Lattanzi, F. A., Rebuffo, M., & Monza, J. (2019). Selection of competitive and efficient rhizobia strains for white clover. Frontiers in Microbiology, 10. 10.3389/fmicb.2019.00768

Lesins, K. A., & Lesins, I. (1979). Genus medicago (leguminosae), a taxogenetic study. VEGETATIO, 50(2), 92–92. 10.1007/BF00055206

Li, H., Zhou, Y., Xin, W., Wei, Y., Zhang, J., & Guo, L. (2019). Wheat breeding in northern China: Achievements and technical advances. The Crop Journal, 7(6), 718–729. 10.1016/j.cj.2019.09.003

Li, X., Zhang, X., Zhao, Q., & Liao, H. (2023). Genetic improvement of legume roots for adaption to acid soils. The Crop Journal, 11(4), 1022–1033. 10.1016/j.cj.2023.04.002

Liu, K., He, R., He, X., Tan, J., Chen, Y., Li, Y., Liu, R., Huang, Y., & Liu, H. (2022). Speed Breeding Scheme of Hot Pepper through Light Environment Modification. Sustainability, 14(19), 12225. 10.3390/su141912225

Luo, D., Zhang, X., Liu, J., Wu, Y., Zhou, Q., Fang, L., & Liu, Z. (2023). DROUGHT-INDUCED UNKNOWN PROTEIN 1 positively modulates drought tolerance in cultivated alfalfa (Medicago sativa L.). The Crop Journal, 11(1), 57–70. 10.1016/j.cj.2022.05.013

Mobini, S. H., & Warkentin, T. D. (2016). A simple and efficient method of in vivo rapid generation technology in pea (Pisum sativum L.). In Vitro Cellular & Developmental Biology - Plant, 52(5), 530–536. 10.1007/s11627-016-9772-7

Mobini, S., Khazaei, H., Warkentin, T. D., & Vandenberg, A. (2020). Shortening the generation cycle in faba bean (Vicia faba) by application of cytokinin and cold stress to assist speed breeding. Plant Breeding, 139(6), 1181–1189. 10.1111/pbr.12868

Narina, S. S., Pathak, S. C., & Bhardwaj, H. L. (2014). Chlorophyll Fluorescence to Evaluate Pigeonpea Breeding Lines and Mungbean for Drought Tolerance. Journal of Agricultural Science, 6(11). 10.5539/jas.v6n11p238

Parajuli, A., Yu, L.-X., Peel, M., See, D., Wagner, S., Norberg, S., & Zhang, Z. (2021). Self-incompatibility, Inbreeding Depression, and Potential to Develop Inbred Lines in Alfalfa (pp. 255–269). 10.1007/978-3-030-74466-3_15

Potts, J., Jangra, S., Michael, V. N., & Wu, X. (2023). Speed Breeding for Crop Improvement and Food Security. Crops, 3(4), 276–291. 10.3390/crops3040025

Putnam, D. H. (2021). Factors Influencing Yield and Quality in Alfalfa (pp. 13–27). 10.1007/978-3-030-74466-3_2

R Core Team ((Version 4.3.2)). (2023). R: A language and environment for statistical computing. R Foundation for Statistical Computing, Vienna, Austria. https://www.R-project.org/

Samac, D. A., & Temple, S. J. (2021). Biotechnology Advances in Alfalfa (pp. 65–86). 10.1007/978-3-030-74466-3_5

Samantara, K., Bohra, A., Mohapatra, S. R., Prihatini, R., Asibe, F., Singh, L., Reyes, V. P., Tiwari, A., Maurya, A. K., Croser, J. S., Wani, S. H., Siddique, K. H. M., & Varshney, R. K. (2022). Breeding More Crops in Less Time: A Perspective on Speed Breeding. Biology, 11(2), 275. 10.3390/biology11020275

Samineni, S., Sen, M., Sajja, S. B., & Gaur, P. M. (2020). Rapid generation advance (RGA) in chickpea to produce up to seven generations per year and enable speed breeding. The Crop Journal, 8(1), 164–169. 10.1016/j.cj.2019.08.003

Sánchez-Reinoso, A. D., Ligarreto-Moreno, G. A., & Restrepo-Díaz, H. (2019). Chlorophyll α Fluorescence Parameters as an Indicator to Identify Drought Susceptibility in Common Bush Bean. Agronomy, 9(9), 526. 10.3390/agronomy9090526

Signorelli, S., Casaretto, E., Sainz, M., Díaz, P., Monza, J., & Borsani, O. (2013). Antioxidant and photosystem II responses contribute to explain the drought–heat contrasting tolerance of two forage legumes. Plant Physiology and Biochemistry, 70, 195–203. 10.1016/j.plaphy.2013.05.028

Suttie, J. (2012). Alfalfa and Relatives: Evolution and Classification of Medicago. By E. Small. Wallingford, UK: CABI (2011), pp. 727, £110.00. ISBN 978-1-84593-750-8. Experimental Agriculture, 48(2), 308–309. 10.1017/S0014479711001384

Sysoeva, M. I., Markovskaya, E. F., & Shibaeva, T. G. (2010). Plants under Continuous Light: A Review. Global Science Books, Plant Stress.

Temesgen, B. (2022). Speed breeding to accelerate crop improvement. International Journal of Agricultural Science and Food Technology, 8(2), 178–186. 10.17352/2455-815X.000161

Vincent, J. M. (1970). A manual for the practical study of the root-nodule bacteria. (Vol. 15).

Watson, A., Ghosh, S., Williams, M. J., Cuddy, W. S., Simmonds, J., Rey, M.-D., Asyraf Md Hatta, M., Hinchliffe, A., Steed, A., Reynolds, D., Adamski, N. M., Breakspear, A., Korolev, A., Rayner, T., Dixon, L. E., Riaz, A., Martin, W., Ryan, M., Edwards, D., … Hickey, L. T. (2018). Speed breeding is a powerful tool to accelerate crop research and breeding. Nature Plants, 4(1), 23–29. 10.1038/s41477-017-0083-8

Wei, J., Fan, Z., Hu, F., Mao, S., Yin, F., Wang, Q., Chai, Q., & Yin, W. (2024). Legume green manure can intensify the function of chemical nitrogen fertilizer substitution via increasing nitrogen supply and uptake of wheat. The Crop Journal, 12(4), 1222–1232. 10.1016/j.cj.2024.07.004

Wickham, H., Chang, W., Henry, L., Pedersen, T. L., Takahashi, K., Wilke, C., Woo, K., Yutani, H., Dunnington, D., & RStudio. (2023). ggplot2: Create elegant data visualisations using the grammar of graphics. (3.3.3). https://ggplot2.tidyverse.org

Wolter, F., Schindele, P., & Puchta, H. (2019). Plant breeding at the speed of light: the power of CRISPR/Cas to generate directed genetic diversity at multiple sites. BMC Plant Biology, 19(1), 176. 10.1186/s12870-019-1775-1

Yi, L.-X., & Kole, C. (2021). The Alfalfa Genome (L.-X. Yu & C. Kole, Eds.). Springer International Publishing. 10.1007/978-3-030-74466-3

Zhao, H., Zhao, S., Cao, Y., Jiang, X., Zhao, L., Li, Z., Wang, M., Yang, R., Zhou, C., Wang, Z., Yuan, F., Ma, D., Lin, H., Liu, W., & Fu, C. (2024). Development of a single transcript CRISPR/Cas9 toolkit for efficient genome editing in autotetraploid alfalfa. The Crop Journal, 12(3), 788–795. 10.1016/j.cj.2024.04.001

Zheng, L., Wen, J., Liu, J., Meng, X., Liu, P., Cao, N., Dong, J., & Wang, T. (2022). From model to alfalfa: Gene editing to obtain semidwarf and prostrate growth habits. The Crop Journal, 10(4), 932–941. 10.1016/j.cj.2021.11.008

